## Supplementary figures and images for "Identification of potential molecular mimicry in pathogen-host interactions"

### SOM Figure S1

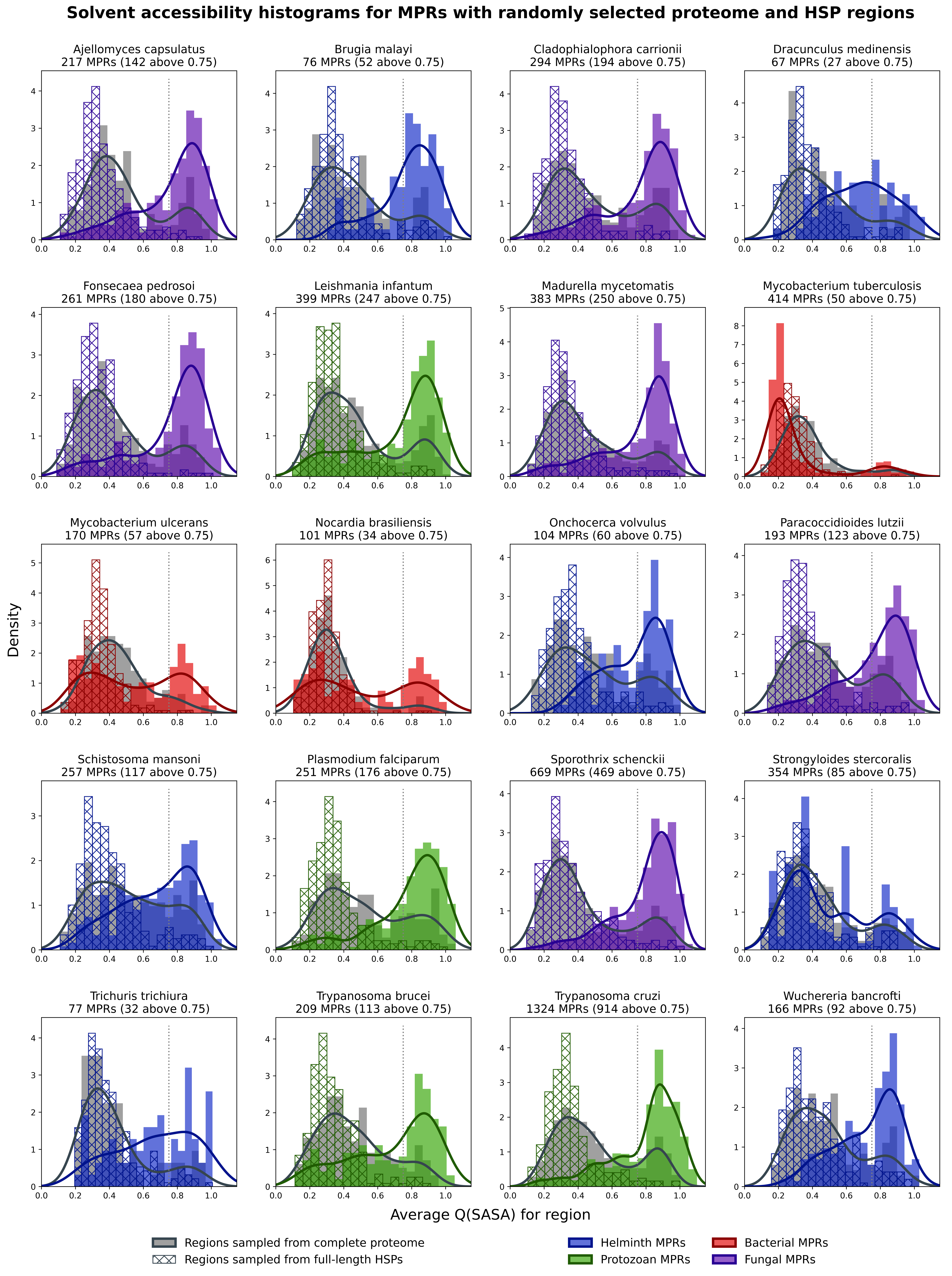

### SOM Figure S2

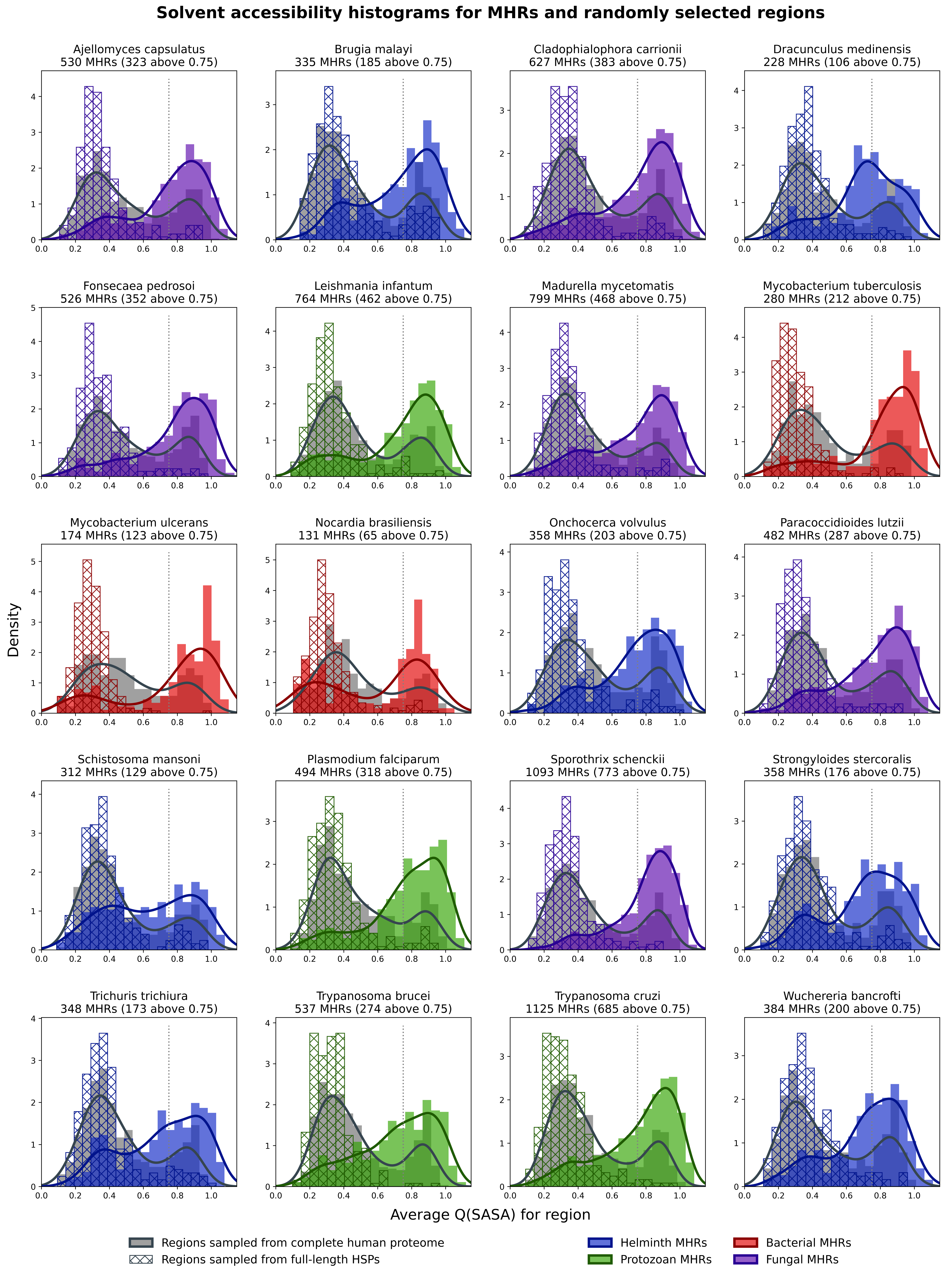
